# Amyloid Beta Oligomers Prevents Lysosomal Targeting of miRNP to Stop Its Recycling and Target Cytokine Repression in Glial Cells

**DOI:** 10.1101/2020.12.24.424324

**Authors:** Dipayan De, Suvendra N. Bhattacharyya

**Affiliations:** RNA Biology Research Laboratory, Molecular Genetics Division, CSIR-Indian Institute of Chemical Biology, Kolkata, India

**Keywords:** miRNA mediated translation repression, mRNA compartmentalization, RNA processing bodies, miRNP recycling, Endosome Lysosome interaction

## Abstract

mRNAs encoding inflammatory cytokines are targeted by miRNAs and remain repressed in neuroglial cells. On exposure to amyloid beta 1-42 oligomers, glial cells start expressing proinflammatory cytokines although there has been increase in repressive miRNAs levels as well. Exploring the mechanism of this potential immunity of target cytokine mRNAs against repressive miRNAs in amyloid beta exposed glial cells, we have identified differential compartmentalization of repressive miRNAs in glial cells to explain this aberrant miRNA function. While the target mRNAs were found to be associated with polysomes attached to endoplasmic reticulum, the miRNPs found to be present predominantly with endosomes that failed to recycle to endoplasmic reticulum attached polysomes to repress mRNA targets in treated cells. Amyloid beta oligomers, by masking the Rab7 proteins on endosomal surface, affects Rab7 interaction with Rab Interacting Lysosomal Protein (RILP) on lysosomes to restrict endosomal maturation and its lysosomal targeting. This causes retarded miRNP targeting to lysosomes and recycling. Similar defects in miRNP retrieval has been observed in endosome maturation defective cells depleted for RILP or treated with Bafilomycin. RNA processing body localization of the miRNPs was also noted in treated cells that happens as a consequence of enhanced endosomal retention of miRNPs. Interestingly, depletion of P-body partly rescues the miRNA function in glial cells exposed to amyloid beta and restricts the excess cytokine expression there.

**Graphical Abstract:** 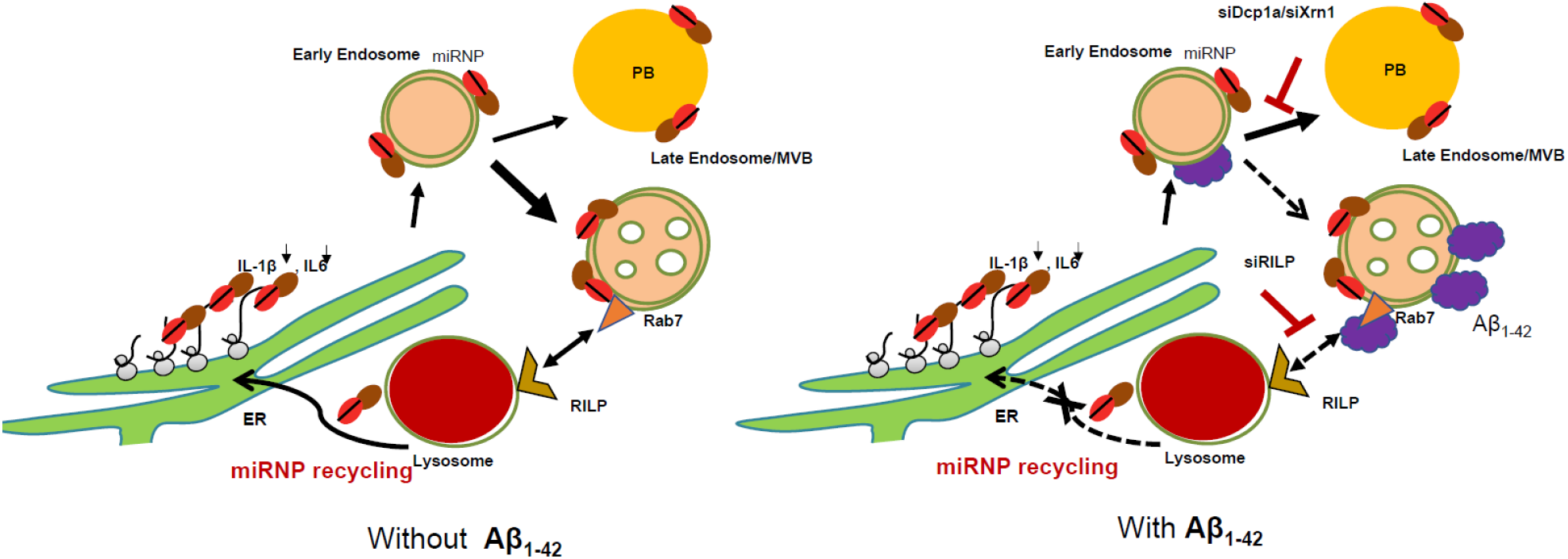

**Key Points:**

- Amyloid beta exposure causes accumulation of inactive miR-146 miRNP to cause elevated proinflammatory cytokine production in glial cells.
- Amyloid beta masks Rab7-RILP interaction to reduce endosome lysosome interaction.
- Accumulated miRNPs failed to get targeted to lysosomes in amyloid exposed cells due to loss of endosome lysosome interaction
- Lysosomal compartmentalization of miRNPs is required for its recycling and repression of *de novo* targets
- Accumulated miRNPs are stored in P-Bodies and depletion of P-Bodies rescues miRNA function in amyloid exposed glial cells.

## Introduction

MicroRNAs (miRNA) are 20-22 nucleotides long non-coding RNAs that represses target mRNAs in both plant and animal cells. miRNAs associate with different Ago proteins (Ago1-4) and form miRNA induced silencing complexes (miRISC) which bind with target mRNAs and either represses or degrades them to stop protein expression (Filipowicz et al. 2008). On average miRNAs have a half-life of 11.4 hrs which is almost 4 times that of mRNAs and thus miRNAs are in general more stable than mRNAs (Reichholf et al. 2019). High stability of miRNA is possibly required to avoid the complicated multistep procedure of miRNA biogenesis and metazoan cells uses each copies of miRNA repeatedly for repression of different target messages instead of making each copies of miRNA for each target mRNAs to repress. Interestingly, at steady state, there are several copies of each miRNAs present in a cell to fine tune a particular cellular event.

miRNA-mediated repression and target degradation are two spatio-temporally uncoupled processes in human cells and the RNA degradation event that happens on the late endosomal membrane plays the pivotal role in recycling of miRNPs and thus for the next round of repression of target mRNAs in mammalian cells by the respective miRNAs (Bose et al. 2017). In this context endoplasmic reticulum (ER) membrane acts as the site where nucleation of miRNA or siRNA take place before the miRNPs encounter with target mRNA (Li et al. 2013; Stalder et al. 2013; Barman and Bhattacharyya 2015; Bose et al. 2020). While perturbation of endosomal trafficking can affect siRNA function (Lee et al. 2009), entrapment of miRNPs with ER restricts its export and abundance (Chakrabarty and Bhattacharyya 2017).

RNA processing bodies are considered as the subcellular structures responsible for reversible storage of miRNA-repressed messages and depletion of P-bodies (PBs) is associated with relocalization of the repressed messages to translating polysomes and thus considered as the key mechanism for reversible regulation of repressive activity of miRNAs (Bhattacharyya et al. 2006). In neuronal context, storage of Ago2 to PBs is considered as the prerequisite for reversal of miRNA activity during growth factor withdrawn and essential for neuronal survival (Patranabis and Bhattacharyya 2018).

In this study, we report how the exposure of amyloid-β oligomers (Aβ_1-42_) of rat glioblastoma cells decreases the cellular miRNA activity by enhancing the sequestration of miRNP complexes to early endosome and PBs. This also causes uncoupling of cytokine mRNAs like IL-1β, IL-6 from Ago2 which accounts for their higher expression in diseased cells. Aβ_1-42_ oligomer perturbs late endosome maturation and its subsequent fusion with lysosomes which is found to be important for miRNP recycling and its binding with newly formed target mRNAs. We propose this mechanism, that explains the increased proinflammatory cytokine expression and neuroinflammation due to defects in miRNA recycling and target mRNA repression in Aβ_1-42_ exposed glial cells.

## Results

### miRNA-derepressed mRNAs are associated with the polysomes in Aβ_1-42_ exposed cell

A 24hrs of Aβ_1-42_ treatment increases mRNA levels of pro-inflammatory cytokines like IL-1β and IL-6 in C6 glioblastoma cells (Fig. 1A). Expression of cytokines are regulated by several miRNAs in mammalian cells, for example IL-6 is a direct target of miRNA let-7a (Iliopoulos et al. 2009). Does the repressive activity of miRNAs get impaired in treated cells? To check the activity of miRNAs upon Aβ_1-42_ exposure, C6 glioma cells were transfected with Renilla Luciferase (RL) reporter having three imperfect miRNA let-7a binding sites (Fig. 1B). Almost a three folds reduction in let-7a activity was observed in Aβ_1-42_ treated cells compared to the DMSO treated control cells (Fig. 1C). However, the total cellular level of let-7a was found to be up regulated upon exposure to Aβ_1-42_ (Fig. 1D). An exogenously expressed liver specific miR-122 in C6 cells also showed similar reduction in repressive activity and increased cellular miRNA level indicating a reduced miRNA activity is not a miRNA identity specific event upon Aβ_1-42_ treatment. The expression of miR-122 was ensured from a pre-miR-122 encoding plasmid having a U6 promoter and change in miR-122 levels in affected cells was normalized again U6 RNA. The increase of only mature miR-122 and not pre-miR-122 level upon Aβ_1-42_ treatment suggests that the increase of miRNA level happens primarily at post transcriptional level and not due to a transcriptional surge of pre-miR-122 in treated cells (Fig. 1D and 1E). As Aβ_1-42_ treatment did not show any decrease in either miRNA or Ago2 protein level, we hypothesized a lack of interaction between miRNA and Ago2 to explain the reduced miRNA activity in amyloid exposed cells. To verify the hypothesis, we used C6 glioma cells expressing FLAG-HA tagged Ago2 (FH-Ago2) and treated them with DMSO or Aβ_1-42_ for 24hrs. FH-Ago2 was immunoprecipitated (IP) using anti-FLAG beads. It was followed by qPCR based measurement of Ago2 bound miRNAs. The qPCR data was normalized against the amount of Ago2 immunoprecipitated from each of the samples. qPCR data revealed an increased association between Ago2 with miR-146a and miR-155 (Fig. 1F), two key miRNAs in macrophage cells which are important regulators of pro-inflammatory cytokines (Sheedy and O’Neill 2008; Kurowska-Stolarska et al. 2011). Contrary to high association of miRNA with Ago2, the interaction between the Ago2 and the target mRNAs was found to be decreased in treated cells as Ago2-association of cytokine IL-1β and IL-6 mRNAs were reduced considerably upon Aβ_1-42_ treatment (Fig. 1G). This suggests the ineffectiveness of the miRNPs accumulated in Aβ_1-42_ treated cells.

**Figure 1.**
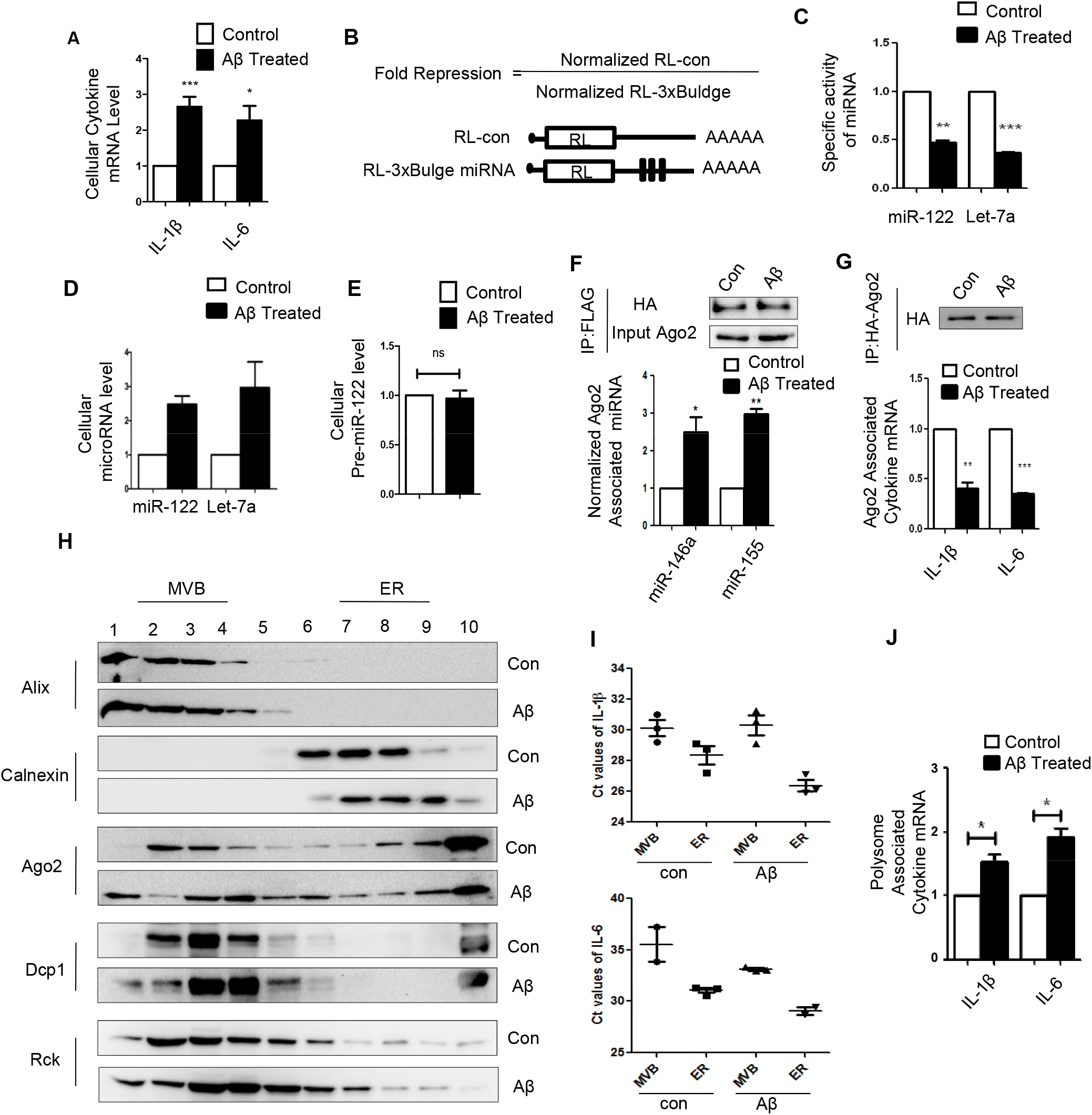
Uncoupling of target mRNA with miRNPs in Aβ_1-42_ exposed cells. **A** qPCR data showing cellular IL-1β and IL-6 level of C6 glioblastoma cell activated with 2.5μM Aβ_1-42_ oligomer for 24h. qPCR data was normalized with GAPDH mRNA level. **B** Schematic design of different Renilla reporters (RL) used for scoring miRNA activity. Respective miRNA binding sites were cloned on 3’UTR of the Renilla mRNA having a perfect and three bulge sites for that miRNA. Fold repression is the ratio of normalized reporter expression vs normalized control. **C,D** Effect of Aβ_1-42_ treatment on cellular miRNA activity and level. Specific activity of each miRNA was calculated by normalizing the fold repression of target reporter mRNA with cellular miRNA level. The repression data was obtained by luciferase based quantification (C), whereas expression level was measured using qPCR and was normalized against the expression of U6 SnRNA (D). **E** Levels of pre-miRNA in cells treated with Aβ_1-42_ oligomers, value normalized against 18sRNA. **F-G** Aβ_1-42_ treated cells show increased Ago2 miRNA association but decreased Ago2 associated cytokine mRNA levels. Immunoprecipitation (IP) of FH-Ago2 from Control and 24h of Aβ_1-42_ (2.5μM) oligomer treated C6 glioblastoma cells expressing FH-Ago2 were done and qRT-PCR based quantification was done. Value normalized to the amount of Ago2 pulled down by immunoprecipitation for miRNA (F) and mRNA (G). **H** Distribution of cytokine mRNA along with Ago2 and few other relevant proteins with endosomal and endoplasmic reticulum fractions in cells treated with Aβ_1-42_. In the upper panel, C6 Glioblastoma cells were activated with 2.5 μM Aβ_1-42_ for 24h before lysis and lysate was ultracentrifuged on a 3-30% iodixanol (optiprep) gradient. Fraction No. 2-4 was designated as MVB/Endosomes and 7-9 was designated as ER. Alix and calnexin were used as markers of endosomes/MVB and endoplasmic reticulum respectively. **I** Distribution of cytokine mRNAs in different subcellular fractions after Aβ_1-42_ treatment in C6 Glioblastoma cells. The mean threshold cycle (Ct) values were plotted for IL-1β and IL-6 mRNA level to check the distribution of cytokine mRNAs in both MVB and ER fractions in control and treated cells. **J** Relative levels of IL-1β and IL-6 mRNA associated with polysomal fraction isolated from control and Aβ_1-42_ treated C6 Glioblastoma cells. The qPCR data was normalized against the GAPDH mRNA present in the polysome fraction. For statistical significance, minimum three independent experiments were considered in each case unless otherwise mentioned and error bars are represented as mean ± S.E.M. P‐values were calculated by utilizing Student’s t-test. ns: non‐significant, *P < 0.05, **P < 0.01, ***P < 0.0001.

For an effective miRNA mediated repression process in mammalian system, all three main components of miRNA-mediated repression i.e. Ago2, miRNA and target mRNA are required to nucleate between them before effective repression could occur. From our previous observations, there was a loss of miRNA activity caused by interaction loss between miRNP and target mRNA. This possible de-linkage of miRNP and target mRNA may happen due to differential localization of miRNP and target messages in distinct sub-cellular compartments. To investigate the sub-cellular localization of miRNP and their target mRNAs, a 3-30% Iodixanol gradient ultracentrifugation was done with cell homogenates obtained from control and Aβ_1-42_ treated C6 cells to separate different sub-cellular organelles. There has been an enrichment of Ago2 in the lighter fractions along with the endosome marker Alix in Aβ_1-42_ treated cells compared to the control cells (Fig. 1H). RNA extraction followed by qPCR based quantification from both Alix (Endosome) and Calnexin enriched (ER) fractions showed higher cytokine mRNAs like IL-1β and IL-6 in ER enriched fractions in Aβ_1-42_ treated cells (Fig. 1I). More specifically, increased association of IL-1β and IL-6 mRNAs was found with polysomal fractions that explain the impaired repression of these mRNAs by miRNAs that leads to increased expression in Aβ_1-42_ treated C6 glioblastoma cells (Fig. 1J).

### Compartmentalization of miRNPs to the Early Endosomes in Aβ_1-42_ oligomers treated glial cells

Our previous experiments revealed Aβ_1-42_ causes reduced interaction between Ago2 and cytokine mRNAs, where derepressed mRNAs are associated with the ER while Ago2 gets enriched in endosomal fractions (Fig. 1G and 1H). There have been previous studies that suggest lysosome targeting of endosome is required for its maturation and that may not happen in isolation (Eden et al. 2010). Endosomes forms contact sites with ER that helps in endosome maturation as well as exchange of cargo between endosome and ER (Rocha et al. 2009). We have checked the possibility whether Aβ_1-42_ treatment could perturb interaction between ER-Endosome to lead to compartmentalization of miRNPs in treated cells. Confocal images were taken for both control and Aβ_1-42_ treated cells, where C6 glioblastoma cells were transiently transfected with YFP-Endo and ER-DsRed which stained early endosome (EE) and ER respectively, whereas Late endosome (LE) were stained with Rab7 antibody (Fig. 2A, S1A). We found a significant decrease in colocalization between both ER-EE and ER-LE pairs upon Aβ_1-42_ treatment (Fig. 2B). LEs are known to interact more with ER than EEs, and EEs shows increased interaction with ER as endosome matures (Friedman et al. 2013). Interestingly, Aβ_1-42_ exposure diminishes the colocalization between ER-EE and ER-LE (Fig. 2B), which may affect endosome maturation process as well. However no such interaction loss between lysosomes and ER were observed in Aβ_1-42_ treated cells (Figure 2B). To confirm the role of Aβ_1-_ 42 in regulating the interaction between ER and EE, an *in vitro* reaction was carried out with enriched EE and ER fraction in presence of Aβ_1-42_ and ATP. An immunoprecipitation of EEA1 followed by western blot revealed reduced interaction between EE and ER upon amyloid exposure (Fig. S1B). To find out the cellular localization of non-functional miRNPs, a 3-15% iodixanol gradient ultracentrifugation was carried out with cell lysates taken from both the control and Aβ_1-42_ treated cells. The early endosome (EE) and late endosomes (LE) were separated on the 3-15% gradient and presence of Ago2 in EE fraction was noted (Fig. 2C and 2D). RNA extraction followed by qPCR based quantification from both HRS (EE) and Rab7 enriched (LE) fractions showed abundance of both miR-146a and miR-155 in EE fractions in Aβ_1-42_ treated cells (Fig. 2E). Enrichment of Ago2 associated miRNA level in EE fraction in Aβ_1-42_ treated cells was also noted (Fig. 2F).

**Figure 2.**
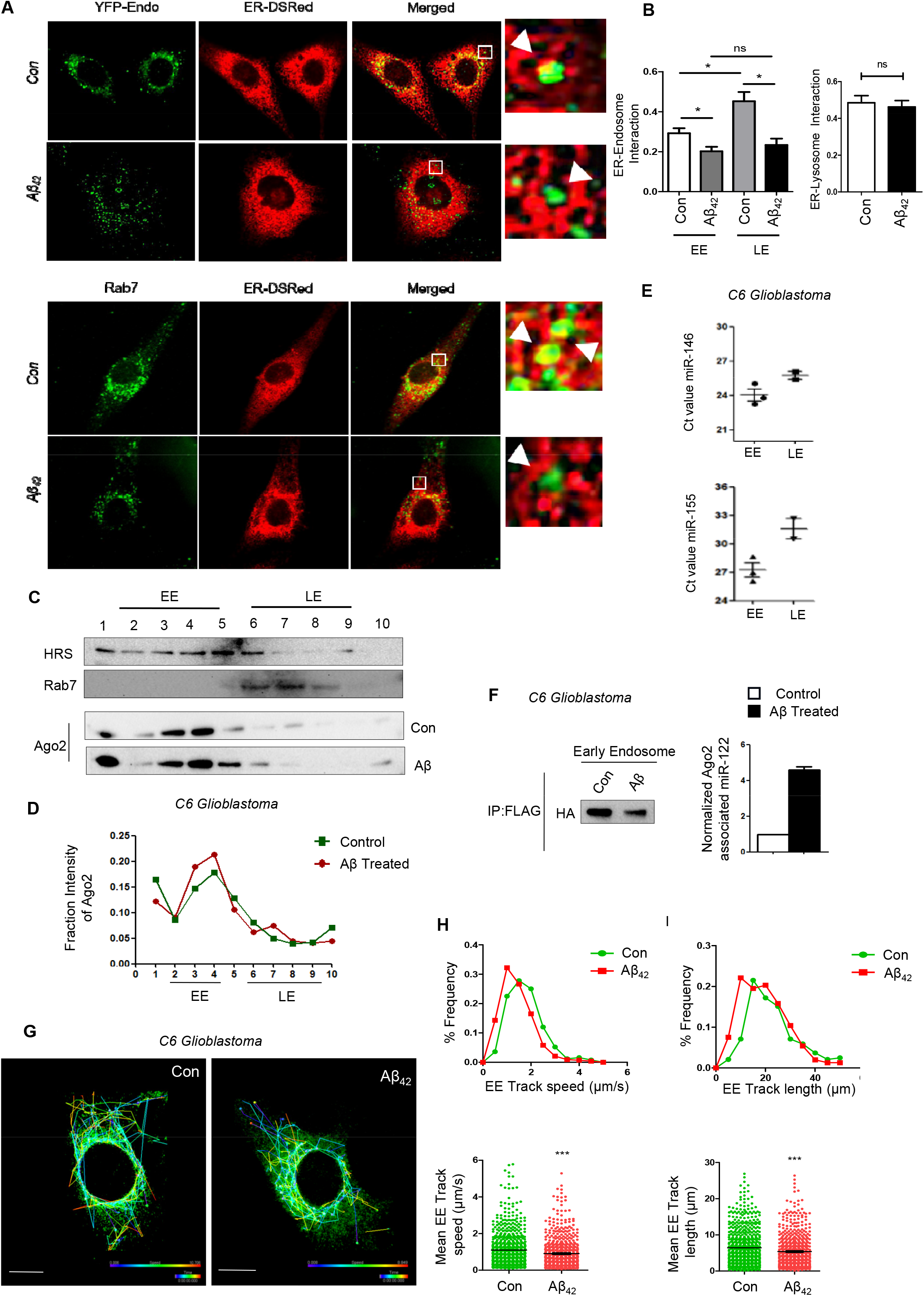
Loss of ER-Endosome interaction retains miRNPs with the Early Endosome. **A-B** Decreased ER-Endosome interaction upon Aβ_1-42_ treatment. Early and Late endosomes were tagged with YFP-Endo and Rab7 respectively. ER was tagged with ER-DsRed. Confocal images showing colocalized area (Yellow) between early and late endosome with ER. Pearson’s coefficient of colocalization was measured for both DMSO and 2.5μM Aβ_1-42_ treatment. Similarly interaction between Lysosomes and ER were also measured and plotted for control and Aβ_1-42_ treated cells (B, right panel). **C-D** Distribution of Ago2 and miRNA on a 3-15% iodixanol (optiprep) gradient to separate the early (EE) and late (LE) endosomes. The optiprep fractions were western blotted for HRS, Rab7 and Ago2 (C). Quantification of percent Ago2 present with each fraction were done and data is plotted in the panel D. HRS and Rab7 were used as the markers of EE and LE respectively. **E** Distribution of miR-146 and miR-155 in EE and LE fraction in Aβ_1-42_ treated cells. The mean threshold cycle (Ct) values were plotted for these miRNAs to check the distribution in both EE and LE fractions in control and treated cells. **F** Estimation of Ago2 associated miR-122 level in both EE and LE fraction in Aβ_1-42_ treated C6 Glioblastoma cells. The qPCR data was normalized with the amount of Ago2 pulled down during immunoprecipitation (IP). **G** Representative image showing tracks of EE analysed from time lapsed video microscopy. Colour code depicting track speed of individual EEs. Bar 10 μm. **H-I** Graphs representing the frequency distribution and mean track speed (H) and frequency distribution and mean track length (I) of individual EEs of both control and Aβ_42_ treated glioblastoma cells. For statistical significance, minimum three independent experiments were considered in each case unless otherwise mentioned and error bars are represented as mean ± S.E.M. P‐values were calculated by utilizing Student’s t-test. ns: non‐significant, *P < 0.05, **P < 0.01, ***P < 0.0001.

In the experiments described above, we found non-functional miRNPs to remain with the early endosome fraction in Aβ_1-42_ treated cells while in mammalian cells it is known to get shuttled to late endosomes along with the repressed mRNAs for degradation, miRNP recycling or MVB entrapment of miRNAs for extracellular export (Mukherjee et al. 2016; Bose et al. 2017). Subsequently, we have studied the endosomal maturation process in Aβ_1-42_ treated cells. The number and surface area of both Early Endosomes (EE) and Late Endosomes (LE) were measured. On exposure of Aβ_1-42_, even though early endosomes were found to be enlarged in glial cells, the number of late endosomes were found to be decreased suggesting a possible retardation of maturation of early to late endosomes (Fig. S2A and 2B). We hypothesized Aβ_1-42_ may affect the mobility of enlarged EE resulting in reduced EE-ER association and EE-LE maturation. We compared the percentage frequency distribution of EE track speed from time lapse live cell microscopy (Fig. 2G). Although majority of both control and Aβ_1-42_ treated EE showed track speed of <2.0 μm/sec, Aβ_1-42_ treated EEs were significantly slower (Fig. 2H). Also we found a significant reduction in mean track length of Aβ_1-42_ treated EEs as compared to control EEs (Fig. 2I). Hence from the above data we could conclude that Aβ_1-42_ may cause a potential decrease in mobility of the EEs.

### Perturbation of endosomal maturation in Aβ_1-42_ treated cells causes defect in miRNP recycling and de novo target RNA repression

Defect in endosomal pathway has been previously reported in different neurological disorders such as Alzheimer’s Disease (AD) and in Neimann Picks disease Type C (Nixon 2005; Maxfield 2014). Also there are reports which suggest early endosome as the site where Amyloid Precursor Protein (APP) localizes to form pathogenic amyloid β protein (PMID: 12761223). Researchers have also reported development of AD in Down Syndrome affected patients at very early age of life while onset of defect in the endosomal pathway of the same has been established in murine model of the aforesaid disease ((Cataldo et al. 2000).

Reduced mobility of EE upon Aβ_1-42_ exposure and loss of endosome-ER interaction could play an important role in endosomal cargo delivery to lysosome. To measure the impact of EE dynamics on lysosomal cargo delivery, we observed a slight reduction in track speed and track length of LEs upon amyloid exposure (Fig. S3A-3C). However, a significant decrease in colocalization between late endosome and lysosome has substantiated the idea of an altered endosome maturation process altogether in Aβ_1-42_ treated cells (Fig. 3A and 3B, S3D). A reduced interaction between late endosomal marker Rab7 and lysosomal protein RILP (Rab Interacting Lysosomal Protein) was found in organellar co-immunoprecipitation assay. This confirmed the reduced fusion of late endosomes to lysosomes upon treatment of Aβ_1-42_ (Fig. S2C and S2D). Therefore it could be the endosomal maturation defect that may be linked with miRNP inactivation observed in glial cells.

**Figure 3.**
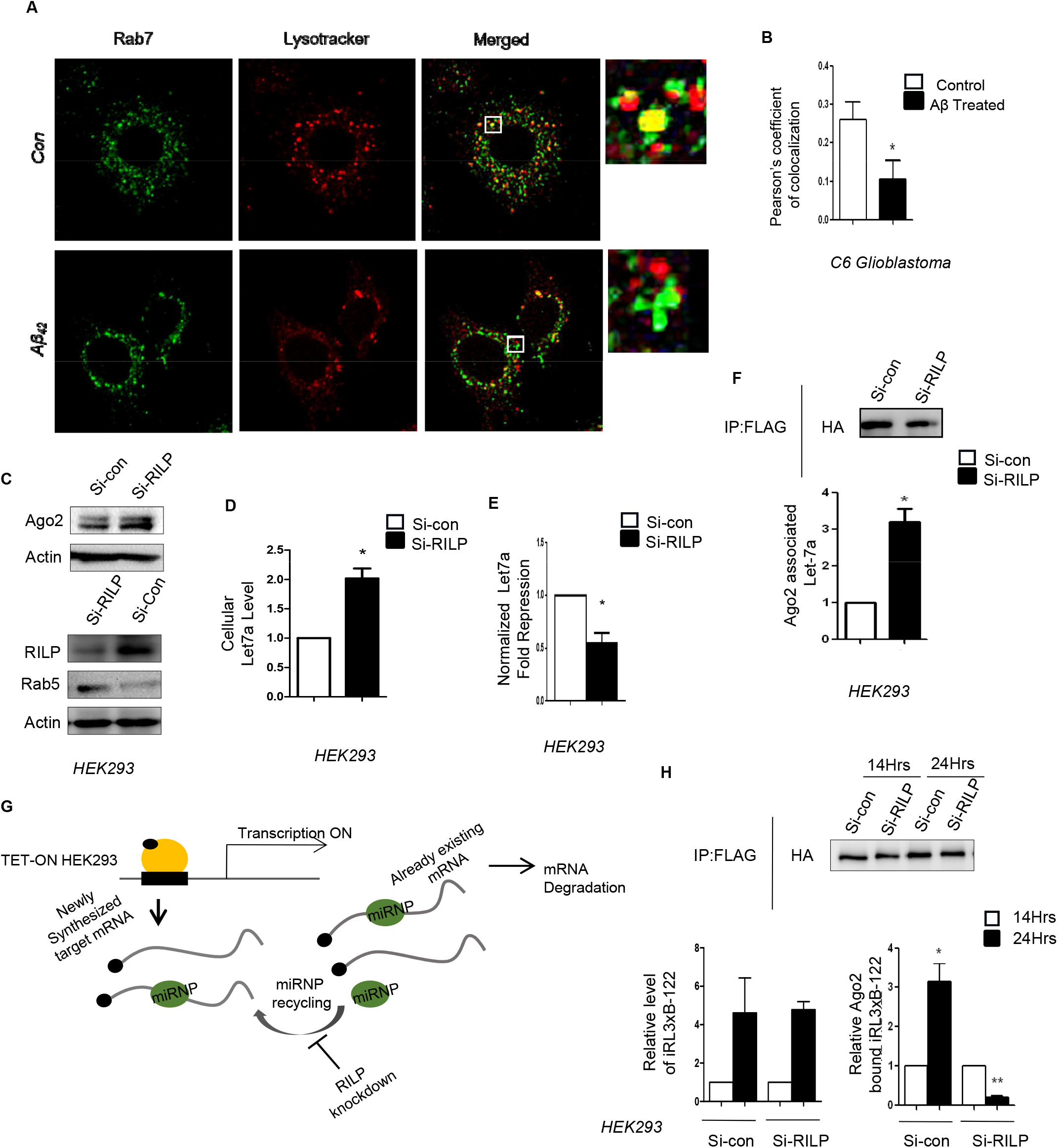
Endosomal maturation is important for miRNA activity regulation and miRNP recycling. **A** Reduced LE-Lysosome interaction upon Aβ_1-42_ exposure in C6 glioblastoma cell. Confocal images showing colocalization between late endosome (LE) and lysosome. LE were marked with Rab7 (green) and lysosomes were stained with lysotracker (Red). Colocalized regions were shown in yellow. **B** Extent of colocalization between late endosome (LE) and lysosome was measured by calculating the Pearson’s coefficient of colocalization between the green(LE) and red (lysosome) pixels. **C** Levels of miRNA interacting and related proteins in HEK293 cells where endosome maturation was blocked. Representative western blot showing the levels of cellular Ago2 in HEK293 cells treated with siRILP or control siRNA (SiCon). β-Actin was used as loading control. Levels of RILP and other endosomal components were also estimated. **D-F** Effect of endosome maturation defect on miRNA levels (D), activity (E) and its Ago2 association (F). Levels of let-7a were measured in cellular, or in Ago2 immunoprecipitated materials in control and siRILP treated cells. RNA was recovered and qRT-PCR estimation was done. The value was normalized against U6 RNA for cellular samples while the amount of immunoprecipitated Ago2 was used for normalization of amount of miRNA let-7a present in the immunoprecipitated materials. Activity of the let-7a was measured by quantifying the repression of a reporter mRNA in control and RILP compromised cells (panel E). **G-H** Defective recycling of miRNPs and poor re-binding to *de novo* synthesized mRNAs in mammalian cells defective for endosome maturation. Control and siRILP treated TET-ON HEK293 cells were co-transfected with inducible RL-3xBulge-miR-122, FLAG-HA-Ago2 and miR-122 expressing pmiR-122 plasmids. After 36h of transfection, RL-3xBulge-miR-122 was induced with Doxycycline for 14h or and 24h a followed by immunoprecipitation of FH-Ago2 to check Ago2 associated reporter mRNA levels. The scheme of the experiments is shown (G). Levels of total and Ago2 associated RL mRNAs were measured and quantified at 14h and 24h post induction (H). RL mRNA levels was normalize with the amount of Ago2 pulled down in the immunoprecipitation experiments. Relative cellular RL mRNAs levels confirm positive induction of reporter target mRNA in both control and RILP knockdown cells. This data was normalized to 18sRNA. For statistical significance, minimum three independent experiments were considered in each case unless otherwise mentioned and error bars are represented as mean ± S.E.M. P‐values were calculated by utilising Student’s t-test. ns: non‐significant, *P < 0.05, **P < 0.01, ***P < 0.0001.

To investigate a possible role of endosomal maturation pathway in miRNA activity regulation, HEK293 cells were knocked down for Rab Interacting Lysosomal Protein (RILP) which is required for the fusion of endosome with lysosome. RILP is a Rab effector protein which facilitates cargo delivery from late endosome to lysosome (Cantalupo et al. 2001; Progida et al. 2007). Knock down of RILP increases both total cellular Ago2 level (Fig. 3C) and let-7a miRNA level (Fig. 3D). To check the effect of downregulation of RILP, on let-7a activity, luciferase assay was done by transfecting a RL-3xbulge-let-7a reporter having three imperfect let-7a binding sites in siRNA treated cells. The assay revealed low miRNA activity when endosome maturation was compromised in HEK293 cells (Fig. 3E). Ago2 pulled down assay revealed increased miRNP levels in RILP compromised cells (Fig. 3F). The data obtained with RILP compromised non-neuronal HEK293 cells have the similar trend of miRNA activity alteration as observed in Aβ_1-42_ treated glioblastoma cells. Additionally, an exposure to Bafilomycin, which blocks endo-lysosomal fusion by inhibiting V-ATPases also increases total Ago2 level along with cellular miRNAs (Fig. S4A and S4B). We also found a reduction in exogenously expressed miR-122 activity (Fig. S4C) and an increase in Ago2 associated miR-122 level in HEK293 cells upon Bafilomycin treatment similar to what has been observed upon RILP knockdown (Fig. S4D). This further establishes the idea that endo-lysosomal fusion has a role in miRNA activity regulation in mammalian cells.

To test further the effect of endosome maturation defect in RILP compromised cells on *de novo* target recognition by miRNPs, miR-122 expressing TET-ON HEK293 cells were used to express miR-122 target RL-3xB-miR122 mRNA in an inducible manner. It was found that Ago2 of RILP knocked down cells showed a reduced association with the *de novo* synthesized mRNA and that may account for the reduced repressive activity we observed in cells depleted for RILP (Fig. 3G and 3H).

### Endosomal retention induced P-body entrapment of miRNP ceases its activity in amyloid beta exposed cells

With expression of a constitutively active GTPase deficient Rab5 mutant, Rab5Q79C, which known to disrupt early to late endosome maturation (Wegner et al. 2010), we documented an excess expression of the protein Dcp1a and RCK/p54 along with Ago2 that are known to get accumulated in P-bodies or PBs in neuronal cells undergoing differentiation (Fig. S5A) (Patranabis and Bhattacharyya 2018). We also noted increased number of PBs in Rab5-CA expressing C6 cells (Fig. S5A-5B). It has been reported previously that endosome maturation and degradation of PB component proteins are linked (Siomi and Siomi 2009). C6 cells when exposed to bafilomycin also showed increased number of Rck/p54 bodies (Fig. S5C-5D). Therefore, with a defect in endosome maturation has its effect on PB components.

Increased aggregation of RNA granules have been reported in different neurological context (Fan and Leung 2016). Re-localization of RNA binding protein TDP43 from nucleus to cytoplasm is a significant phenomenon in Amyotrophic Lateral Sclerosis (ALS) (Chen-Plotkin et al. 2010). Researchers have also reported the presence of Huntington protein (Htt) in PBs which was co-purified with Ago2 protein (Savas et al. 2008).

So the abundance of PBs in Aβ_1-42_ exposed cells could be the direct cause of pathogenesis and that, by inactivating the miRNPs, may account for the reduced miRNA activity but not abundance in the disease context. To investigate the possible role of RNA processing bodies (P-bodies or PBs) in miRNA activity regulation, rat glioblastoma cells were stained for different markers of PBs and confocal imaging was done to measure the size, number of different bodies positive for Dcp1a and Rck/p54. Colocalization between these two proteins was also estimated (Fig. 4A). We found significant increase in Dcp1a positive body size and also increased colocalized Dcp1a and Rck/p54 in PBs after Aβ_1-42_ treatment (Fig. 4B). It was accompanied by almost 2.5 fold increase in both Dcp1a and Rck/p54 body number (Fig. 4C). We also noted approximately a 2 fold increase in colocalization between Dcp1a and Rck/p54 bodies indicating a correlation between PBs size and numbers with Aβ_1-42_ treatment (Fig. 4D). In that context, expression of PBs components also increased in Aβ_1-42_ exposed cells (Fig. 4E). To investigate the possibility of sequestered Ago2 in PBS in Aβ_1-42_ treated cells, C6 glioma cells were transiently transfected with GFP-Ago2 and then counter stained for Rck/p54 a marker of PBs (Fig. 4F). Results showed a significant increase in Ago2 positive bodies (Fig. 4G). We also found a threefold increase in colocalization coefficient between Ago2 positive bodies with Rck/p54 positive bodies indicating that, in Aβ_1-42_ treated cells, endosome-sequestered Ago2 miRNPs translocate to the PBs that may contributes to the reduced miRNP activity observed in treated cells (Fig. 4G).

**Figure 4.**
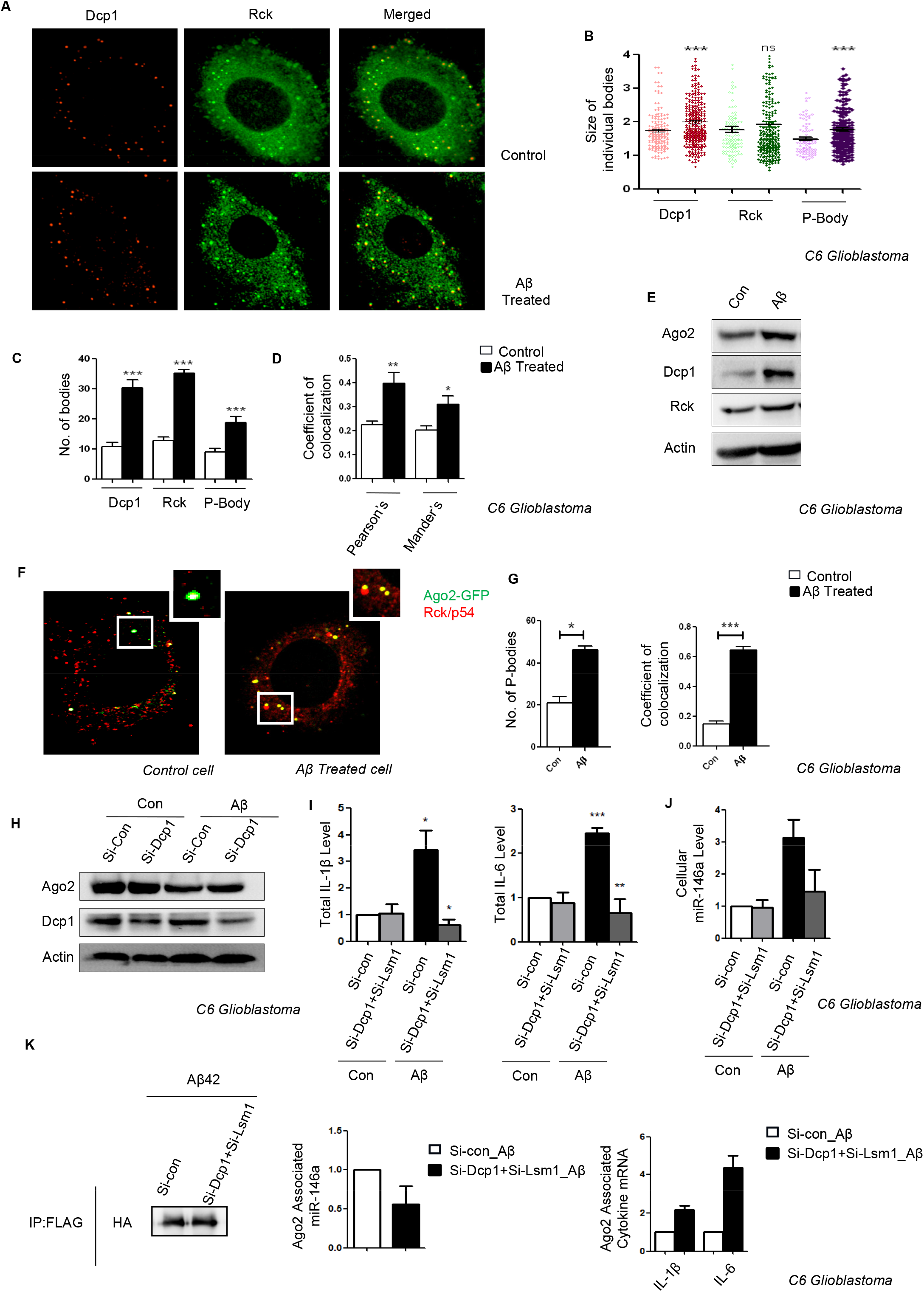
Aβ_1-42_ Activated cells show increase in P-body targeting of Ago2 and deactivation of P-body restores miRNA activity. **A-D** Increased number of P-bodies in cells treated with Aβ_1-42_ oligomers. Colocalization of endogenous Dcp1a (Green) with endogenous Rck/p54 upon 24h of treatment of C6 Glioblastoma cells with 2.5 μM Aβ_1-42_ oligomers (A). Size of the individual Dcp1a, Rck/p54 bodies were measured and plotted for both control and Aβ_1-42_ treated C6 Glioblastoma cells. The plot was generated from 3 individual experiments (mimimum 10 cell/experiment) (B). Number of individual bodies was calculated for each cell for both control and Aβ_1-42_ treated C6 Glioblastoma cells. The plot was generated from 3 individual experiments (minimum 10 cell/experiment) (C). Colocalization of Dcp1 with Rck bodies were measured by calculating Pearson’s and Mander’s co-efficient in control and Aβ_1-42_ treated C6 Glioblastoma cells (D). **E** Levels of expression of PB components in control and Aβ_1-42_ treated C6 Glioblastoma cells. Western blot analysis was done for several protein factors and β-actin was used as loading control. **F-G** Increased Colocalization between Ago2 and Rck/p54 postive bodies. Transiently expressed GFP-Ago2 (green) with the endogenous Rck/p54 (red) in control and Aβ_1-_ 42 treated C6 glioblastoma cells (F). Number of Ago2 bodies per cell was calculated and was plotted for both control and Aβ_1-42_ treated C6 glioblastoma cells. Translocation of Ago2 to the RCK/p54 postive PBs were measure by calculating the pearson’s co-efficient of colocalization between transiently transfected GFP-Ago2 with the endogenous Rck/p54 protein (G). **H-J** Effect of depletion of PB components on Aβ_1-42_ induced miRNA and cytokine expression. Effect of siRNA treatment on expression of Dcp1a in control and siRNA treated cells (H). Relative mRNA level of expression of IL-6 and IL-1β in control and in PB depleted (siDcp1a and siLsm1 treated in combination) both in non-treated and Aβ_1-42_ oligomer treated cells (I). for miR-146 quantification was done by qRT-PCR based estimation. For mRNA GAPDH levels were used for normalization. miR-146 levels were similarly measured (J). U6 snRNA was used for miRNA levels normalization. Values at un-treated control cells were used as unit. **K** Relative cytokine mRNA and miRNA association with Ago2 in Aβ_1-42_ treated control and PB depleted cells. Amount of Ago2 isolated by immunoprecipitation was determined and also used for isolation and quantification of associated mRNA and miRNA levels. qRT-PCR based methods were used for RNA estimation. For statistical significance, minimum three independent experiments were considered in each case unless otherwise mentioned and error bars are represented as mean ± S.E.M. P‐values were calculated by utilising Student’s t-test. ns: non‐significant, *P < 0.05, **P < 0.01, ***P < 0.0001.

How the Aβ_1-42_ oligomers affecting endosome maturation leading to accumulation of miRNPs and their entrapment in PBs is an interesting question. By microscopic analysis we documented proximal localization of Aβ deposition with early endosomes and P-bodies (Fig. S6A-6D). We further noted using STED microscopy, early endosome and P-bodies coming in close contact with each other in Aβ_1-42_ treated cells (Fig. S6E).To make a connection with miRNP-entrapment with PBs and excess cytokine production, we depleted PBs by treating C6 cells by specific siRNAs before exposed them to Aβ_1-42_ oligomer and measured the cytokine content. Cells pre-treated with siRNAs against Dcp1a and Lsm1-two important PB components, showed a reduction in expression of IL-1β and IL-6 compared to siCon transfected cells. This result suggests the importance of miRNP entrapment in PBs with cytokine expression in Aβ treated cells (Fig.4H–4I). In this context the cellular cytokine mRNA and repressive miRNA associated with Ago2 also got reduced, whereas Ago2 associated cytokine mRNA levels increased in PB component depleted cells (Fig. 4J-4K). Our data indicates P-body depletion can potentially decrease the neuroinflammation process by rescuing miRNP recycling defect in Aβ_1-_ 42 treated glioblastoma cells.

## Discussion

Role of different cell organelle in regulating miRNA mediated gene expression in mammalian cells is an interesting but seldom explored area. Membrane and non-membrane bound cell organelles divides the cell cytoplasm into different compartments and this compartmentalization is important in carrying out various biochemical processes such as transcription, translation, protein degradation etc. This compartmentalization also plays a crucial role in fine tuning miRNA activity. Previous reports have highlighted ER membrane as the the site where miRNP nucleation and target mRNA repression take place (Li et al. 2013; Stalder et al. 2013). Our previous work indicates although ER membrane is believed to be the site where target mRNA repression occurs, late endosome serves as the site of mRNA degradation (Bose et al. 2017). In another work it was highlighted how miRNA free Ago2 accumulation affects its MVB to ER translocation and miRNP biogenesis (Bose et al. 2020).

Several neurodegenerative diseases like AD, PD and ALS are prominently linked with EE dysfunction. Enlargement of Rab5 positive vesicles are shown to be one of early symptoms in AD patients (Cataldo et al. 2000). In ALS, problem in GDP/GTP exchange causes hyperactivation of Rab5 leading to early endosome accumulation (Otomo et al. 2003; Lai et al. 2009). How EE dysfunction affects wide variety of diseases? It is possible that EE dysfunction also affects downstream LE maturation and also LE-Lysosome fusion to have its wider implication in health and diseases. Lysosome is one of the most important organelle that removes cellular waste and aberrant proteins to control protein homeostasis in an eukaryotic cell. Niemann-Pick disease type C (NPC) is a pathologic condition involving accumulation of different lipid molecules like sphingolipid and cholesterol in lysosomal lumen (Butler et al. 1993; Vance 2006),. Extensive works have been done on the role of autophagy in protein homeostasis and organelle turnover in neurodegenerative diseases. But one key question remained unanswered is how this EE dysfunction or a problem in endo-lysosomal fusion triggers innate immune system. In this study we have tried to explain how impaired endo-lysosomal fusion leads to accumulation of miRNPs within EE and P-Bodies which fail to recognize the newly formed target mRNA. This mechanism causes a global decrease in miRNA activity and its inability to repress cytokine mRNA in glial cells. Here we have shown importance of cellular organelles which affects the miRNA recycling process suggesting how cellular structures fine tunes the delicate gene expression mechanism during the pathological events.

Do Aβ_1-42_ oligomers have any low-complexity sequence or any short linear motifs required to form any liquid-liquid phase separation? This has not been explored. PBs are example of a separated phase within the cytoplasm where RNA binding proteins along with repressed mRNA can be stored (Filipowicz et al. 2008). Aβ_1-42_ may play a role in increasing local concentration of RNA and RNA binding proteins in cytoplasm to form the PBs which we found frequently in glioblastoma cells exposed to Aβ_1-42_. Researchers made similar observation in Huntington’s disease where Ago2 was found in the Stress Granules (SG) which may account for its inactivation (Pircs et al. 2018). Past investigations also revealed a strong correlation between RNP hyperaggregation with different neurodegenerative diseases like ALS or HD (Savas et al. 2008; Chen-Plotkin et al. 2010) but whether and how these dynamic RNP granules adds to the pathogenesis of these diseases has been an interesting topic to understand. Finally, another important aspect would be to look at whether these phase separated RNP droplets may have other cell biological role to play rather than serve as RNA storage sites. As these droplets are very dynamic in nature and are constantly exchange cargo between soluble cytosol and the insoluble granules, they might have a role to play in different rapid physiological changes such as cell signaling or in ion exchange processes.

Another key aspect would be to look at what is the fate of this accumulated miRNPs. We do know that part of this miRNPs translocate to P-body. This increase in P-body can initiate pathogenesis by inducing some signaling process or by regulating mRNA repression. Another part of this accumulated miRNAs along with miRNPs can be packaged into Extracellular Vesicles (EVs) or exosome and can be exported out of cells (Unpublished Data). This extracellular miRNAs can acts as exocrine signal as they are taken up by recipient glial cells and neurons. The transfer of miRNAs and miRNPs could thus serve as a signal for disease initiation before these cells are actually exposed to β-amyloid. So this process can serve as a protection for the cells which are not exposed to pathogenic proteins. Presence of this exosomes in CSF or in blood can be used for biomarkers for early disease detection as well. It is also noteworthy that HuR protein which can reverse the miRNA mediated repression (Bhattacharyya et al. 2006), is also responsible for extracellular vesicle mediated transfer of miRNAs (Mukherjee et al. 2016) can be targeted and can be a potential therapeutic strategies against these diseases.

### Experimental Procedures

#### Cell culture and transfection

Both C6 glioblastoma and HEK293 cells were cultured in high glucose DMEM medium (Life Technologies) containing 2mM L-glutamine and 10% heat inactivated FCS (Gibco). All the plasmids and si-RNAs were transfected with Lipofectamine-2000 and RNAiMax (Life Technologies) respect9vely following manufacture’s protocol. All the SMARTpool ONTARGETplus si-RNAs were brought from Dharmacon.

TET-ON HEK293 cells were used to carry out experiments using inducible construct. TET-ON HEK293 cells were cultured in DMEM supplemented with 10% TET-approved FCS (Clonetech). Specific genes were induced using 300ng/ml of Doxycycline (Sigma) for desired time points.

#### Preparation of Aβ_1-42_ oligomer

HPLC-purified lyophilized Aβ_1-42_ (American Peptide) was reconstituted in 100% 1,1,1,3,3,3 hexafluoro-2-propanol (HFIP). HFIP was removed by evaporation in SpeedVacand resuspended in 5mM DMSO. The stock was diluted with PBS to 400 mM, and SDS was added to a final concentration of 0.2%. The resulting solution was incubated overnight and diluted again with PBS to 100 mM followed by incubation at 37°C for 18–24 h before use.

#### RNA isolation and Real time PCR

Total RNA was isolated using TRIzol reagent (Life Technologies) following manufacture’s instruction. cDNA was prepared taking 50ng and 200ng of total RNA for miRNA and mRNA respectively. For real time analysis of specific mRNA, cDNA was prepared using random nonamer (Eurogentec reverse transcriptase core kit) followed by real time PCR using Mesa Green quantitative PCR (qPCR) master mix plus (Eurogentec) following manufacture’s protocol. For quantification of specific miRNA,a Taqman reverse transcription kit (Applied Biosystems) was used for cDNA preparation followed by real time PCR using TaqManuniversal PCR mix(Applied Biosystems). qPCR was done using specific taqman based miRNA primers (Supplementry Tables) following manufacture’s instruction. All PCR reactions were done in a 7500 Applied Biosystems real-time system or a Bio-Rad CFX96 real-time system. 18s rRNA and GAPDH mRNA levels were used as a loading control for mRNA quantification whereas U6 snRNA was used loading control for miRNA quantification.

#### Luciferase Assay

Renilla luciferase (RL) and firefly luciferase (FF) activities were measured using a Dual luciferase assay kit (Promega). Cells were transfected with 20ng of RL-Con or RL-3x buldge-Let-7a or RL-3x buldge-miR122 reporter along with 200ng of firefly luciferase. Cells were lysed with 1x Passive lysis buffer (Promega) after 48hrs of transfection and luciferase activity was measured on a VICTOR X3 Plate Reader following manufacture’s protocol. FF normalized RL values were used to score miRNA repression level. Specific activity of miRNAs were calculated normalizing miRNA repression level with total miRNA level.

#### Immunoprecipitation

For immunoprecipitation (IP) cells were lysed with lysis buffer (20 mM Tris-HCl pH 7.5, 150 mM KCl, 5mM MgCl2, 1mM DTT), 0.5% Triton X-100, 0.5% sodium deoxycholate and 1X EDTA-free protease inhibitor cocktail (Roche) for 30 min at 4⁰C followed by three 10sec pulse of sonication. Lysate was cleared at 16000g for 10mins. Protein G agarose beads were blocked in lysis buffer containing 5% BSA for 1hr then incubated with required antibody (final dilution 1:100) for 3hr at 4°C. Cell lysate was incubated with the antibody attached bead for overnight at 4°C. Beads were washed thrice with 1x IP buffer (20 mM Tris-HCl pH 7.5, 150 mM KCl, 5 mM MgCl2, 1mM DTT) and separated into two halves for RNA and protein estimation.

#### Optiprep density gradient centrifugation

For subcellular organelle fractionation, a 3-30% and 3-15% continuous gradient was prepared using OptiprepTM (Sigma-Aldrich, USA) in a buffer constituting 78 mM KCl,4 mM MgCl2,8.4 mM CaCl2, 10 mM EGTA, 50 mM Hepes (pH 7.0). Cells were rinsed with ice cold PBS and a Dounce homogenizer was used to homogenized in a buffer containing 0.25 M sucrose,78 mMKCl,4 mM MgCl2,8.4 mM CaCl2,10 mM EGTA,50 mM Hepes pH 7.0 supplemented with 100μg/ml of Cycloheximide, 5 mM Vanadyl Ribonucleoside Complex (VRC) (Sigma Aldrich), 0.5mM DTT and 1X Protease Inhibitor. The lysate was subjected to centrifugation twice at 1000xg for 5minutes for clarification and placed on top of the prepared gradient. Ultracentrifugation was performed for 5h at 36000 rpm on a SW61T rotor to separate each gradient. 10 fractions were collected by aspiration and further analyzed for RNA and proteins.

#### Polysome isolation

In order to isolate total polysome, buffer constituting 10 mM HEPES pH 8.0, 25 mM KCl, 5 mM MgCl2, 1 mM DTT, 5 mM vanadyl ribonucleoside complex, 1% Triton X-100, 1% sodium deoxycholate and 1 × EDTA-free protease inhibitor cocktail (Roche) supplemented with Cycloheximide (100 μg ml^−1^; Calbiochem) was used to lyse the cells. The clearance of lysate was performed at 3,000X g for 10 min followed by a pre-clearance shift at 20,000X g for 10 min at 4 °C. The loading of the clear lysate was done on a 30% sucrose cushion and ultracentrifuged at 100,000X g for 1 h at 4 °C.The washing of the sucrose cushion was done with a buffer (10 mM HEPES pH 8.0, 25 mM KCl, 5 mM MgCl2, 1 mM DTT), followed by ultracentrifugation for extra 30 min with final resuspension of the pellet in polysome buffer (10 mM HEPES pH 8.0, 25 mM KCl, 5 mM MgCl2, 1 mM DTT, 5 mM vanadyl ribonucleoside complex, 1x EDTA-free protease inhibitor cocktail) for further isolation of RNA and protein from the same.

#### Immunoblotting

Protein samples from whole cell lysate, immunoprecipitated proteins or proteins from cell fractionation were subjected to SDS-PAGE analysis. Western blotting was done on a PVDF membrane overnight at 4°C. Membranes were blocked with 3% BSA for 1hr then probed with specific antibodies (Table No.). Images of western blots were taken with UVP BioImager 600 system equipped with VisionWorks Life Science software, version 6.80 (UVP). Band intensities were calculated using ImageJ software.

#### Immunofluorescence

Cells were grown on 18mm round coverslips and transfected as described earlier. Cell were fixed with 4% paraformaldehyde for 20mins at room temperature in the dark. For indirect immunofluorescence fixed cells were blocked and permeabilized with PBS containing 3% BSA and 0.1% Triton X-100 for 30mins at room temperature. Coverslips were probed with specific antibodies for 16hrs at 4 °C. Coverslips were washed thrice with 1X PBS and then probed with specific AlexaFluor secondary antibody tagged with fluorochrome (dilution 1:500) for 1hr at room temperature.

#### Confocal imaging and post capture image analysis

Confocal fixed cell images were taken with Zeiss LSM800 confocal microscope and analyzed with Imaris7 and ImageJ software. Pearson’s coefficient of colocalization was calculated using Coloc plug-in of Imaris7 software. 3D reconstructions of specific bodies were done using Surpass plug-in of Imaris7 software. Numbers of individual bodies or vesicles were measured using particle generator of Surpass plug-in.

#### Live cell imaging and endosome dynamics

For live cell microscopy cells were transiently transfected with YFP-Endo, GFP-Rab7 or pDsRed2-ER (Clontech). Imaging was done 48hrs after transfection with Leica DMI6000 B inverted microscope equipped with Plan Apo100×/1.40 oil objective (Leica TCS SP8 confocal system). Endosome dynamics was calculated after assigning each vesicle with one particle using surpass plug-in. Endosome track speed and track length were calculated by applying particle tracking algorithm and gap close algorithm.

#### In-vitro organellar interaction

For in-vitro EE-ER interaction study C6 cells were lysed in Dounce homogenizer in a buffer containing 0.25M sucrose,78mM KCl,4 mM MgCl_2_,8.4 mM CaCl_2_,10mM EGTA, 50mM Hepes pH 7.0 supplemented with 100μg/ml of Cycloheximide, 5mM Vanadyl Ribonucleoside Complex (VRC) (Sigma Aldrich), 0.5mM DTT and 1X Protease Inhibitor. Cell lysate was loaded on top of a 3-30% Optiprep gradient as explained earlier and ultra centrifuged at 36000 rpm for 5hrs. Fraction number 2-4 enriched in endosome and 7-9 enriched in ER were further ultra centrifuged for 2hrs at a speed of 50000 rpm and 36000 rpm to isolate endosome and ER respectively. Endosomes were suspended in a buffer containing 0.25M sucrose,78mM KCl,4 mM MgCl_2_,8.4 mM CaCl_2_,10mM EGTA, 50mM Hepes pH 7.0 supplemented with 1X Protease Inhibitor along with 2.5μM Aβ_1-42_ and kept for 1hr at 37°C. ER fraction was added after 1hr along with 1mM ATP and in-vitro interaction was carried out 1hr at 37°C. EE was isolated by immunoprecipitating EE with Protein G-Agarose bead tagged with EEA1 antibody and western blot was done to check amount of Calnexin interacting with EEA1.

#### Statistical analysis

All graphs and statistical analyses were done using Graphpad prism 5.0 (GraphPad, San Diego, CA, USA). Student t-test was done to determine p-value. P value <0.05 was considered as significant. All the experiments were done at least three times. Error bars indicate mean ± S.E.M.

## Acknowledgement

We acknowledge Witold Filipowicz, Gunter Meister and J.M. Backer for different plasmids constructs. Subhas Biswas helped us with primary cells and reagents. SNB is supported by The Swarnajayanti Fellowship from Dept. of Science and Technology, Govt. of India, while D.D. received his support from CSIR, India. We were supported by funds from High Risk High Reward Project Grant, Dept. of Science and Technology, Govt. of India.

## Author contributions

S.N.B. Conceptualize the project, designed research and analyzed data; D.D. performed research and experiments. S.N.B. and D.D. analyzed data and wrote the paper.

The authors declare no competing interest.

---

**Supplementary Figure S1.**
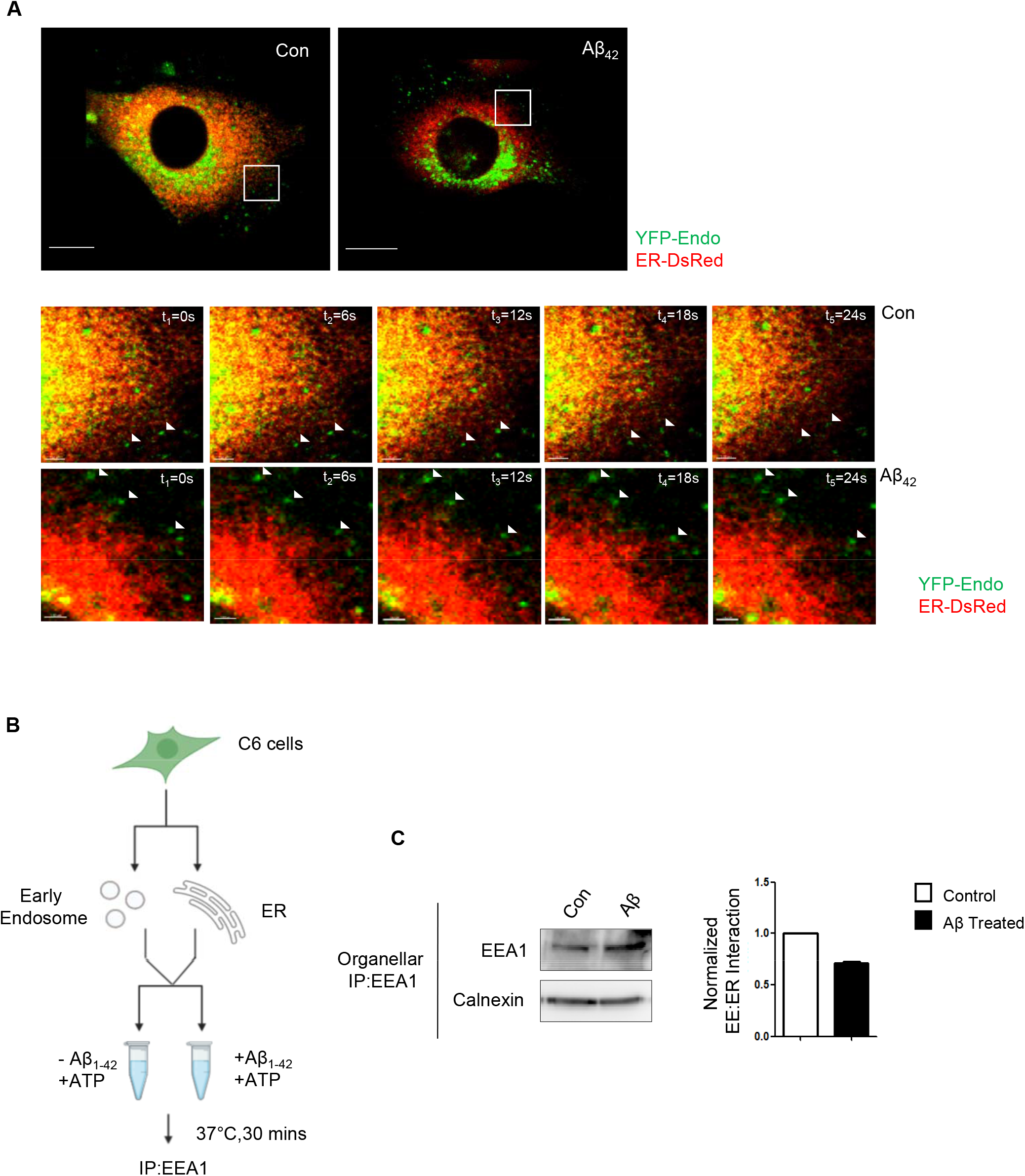
Aβ_1-42_ reduces interaction between EE and ER. **A** Time lapse microscopic images showing interaction between EE (green) and ER (red). C6 glioblastoma cells were transiently transfected with YFP-Endo (green) and ER-DsRed (red) followed by 24 hrs of Aβ_1-42_ treatment. Time lapse images were taken with 6 sec gap for 90 seconds. scale bar 10 μm. **B** Experimental flow-chart to measure the *in vitro* EE-ER interaction change upon Aβ_1-42_ exposure. Both EE and ER were isolated from C6 glioblastoma cells. EEs were pre-incubated with 2.5μM Aβ_1-42_ for 30mins at 37°C followed by further incubation with ER and 1mM ATP for 30mins at 37°C. EEs were immunopurified with anti-EEA1 antibody tagged with protein G-Agarose bead. **C** Western blot showing the amount of calnexin obtained with EEA1 immunoprecipitate ontained from control and treated reasction. Calnexin used as a marker of ER and EEA1 showing the amount of EEs IPed from the *in vitro* reactions. A quantification of amount of calnexin IPed with EEA1 done in triplicate experiments is plotted in right panel. For statistical significance, minimum three independent experiments were considered in each case unless otherwise mentioned and error bars are represented as mean ± S.E.M. P‐values were calculated by utilising Student’s t-test. ns: non‐significant, *P < 0.05, **P < 0.01, ***P < 0.0001.

**Supplementary Figure S2.**
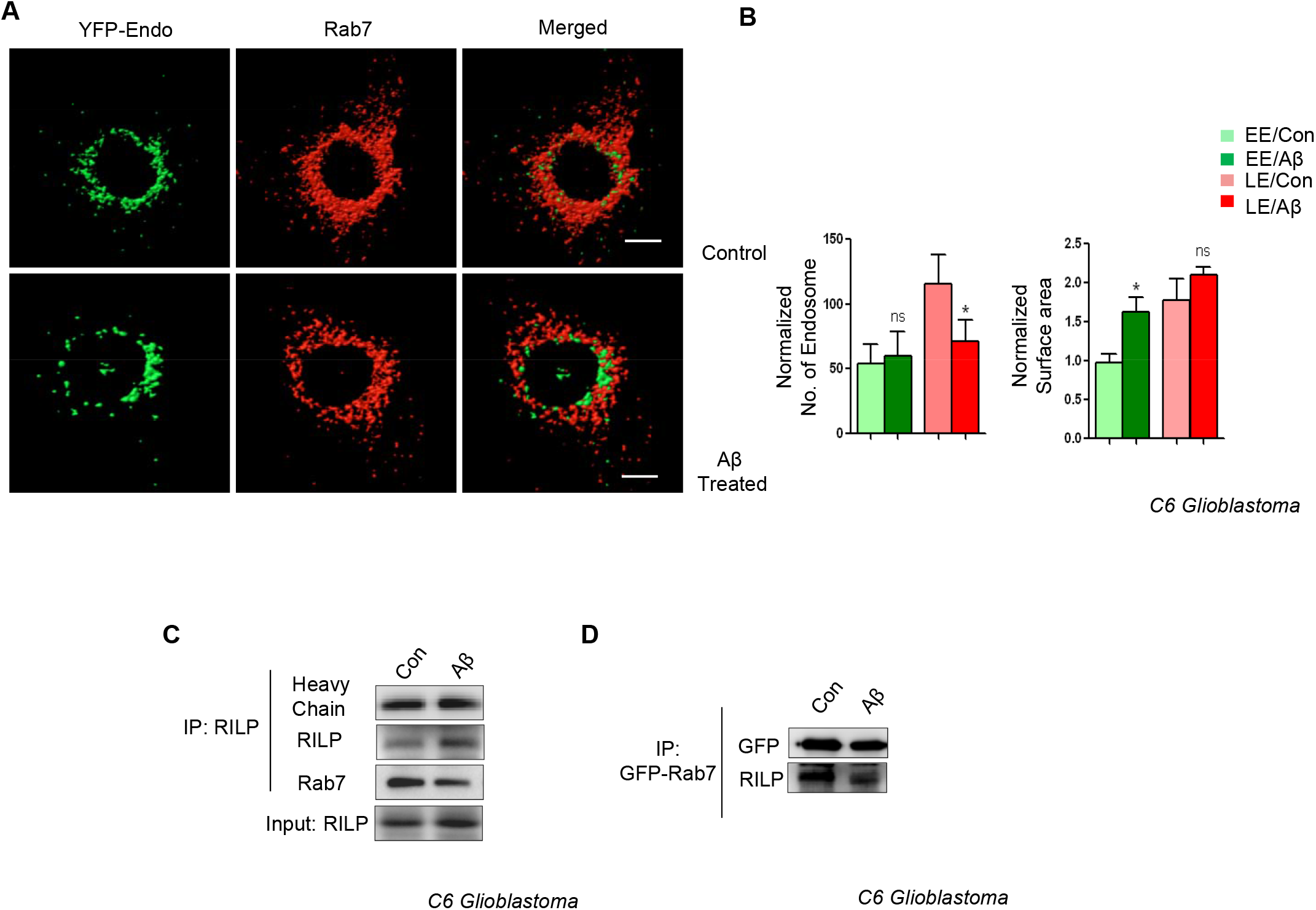
Aβ_1-42_ perturbs endosomal maturation and its subsequent fusion with lysosome. **A-B** Number and size of both early endosome (EE) and late endosome (LE) were calculated for control and Aβ_1-42_ treated C6 Glioblastoma cells by confocal images (A), followed by 3D surface reconstruction. Early endosome (EE) were labeled with YFP-Endo (green) and late endosome (LE) were labelled with endogenous Rab7 (Red). Number of early endosomes (EE) and number of late endosomes (LE) were calculated by counting the number of green vesicles and red vesicles in a specific single cell. The surface area of each of the vesicle was measured and normalized against the surface area of the nucleus. The surface area of the control cell EE was taken as 1 and rest of the data was plotted with respect to it (B). **C** Representative western blot showing amount of Rab7 immunoprecipitated with RILP from control and Aβ_1-42_ treated C6 Glioblastoma cells. Endogenous RILP was pulled down from cells using anti-RILP antibody. **D** Western blot showing amount of RILP immunoprecipitated with Rab7. GFP-Rab 7 was transiently transfected in C6 Glioblastoma cells, followed by IP with GFP antibody from control and Aβ_1-42_ treated C6 Glioblastoma cells.

**Supplementary Figure S3.**
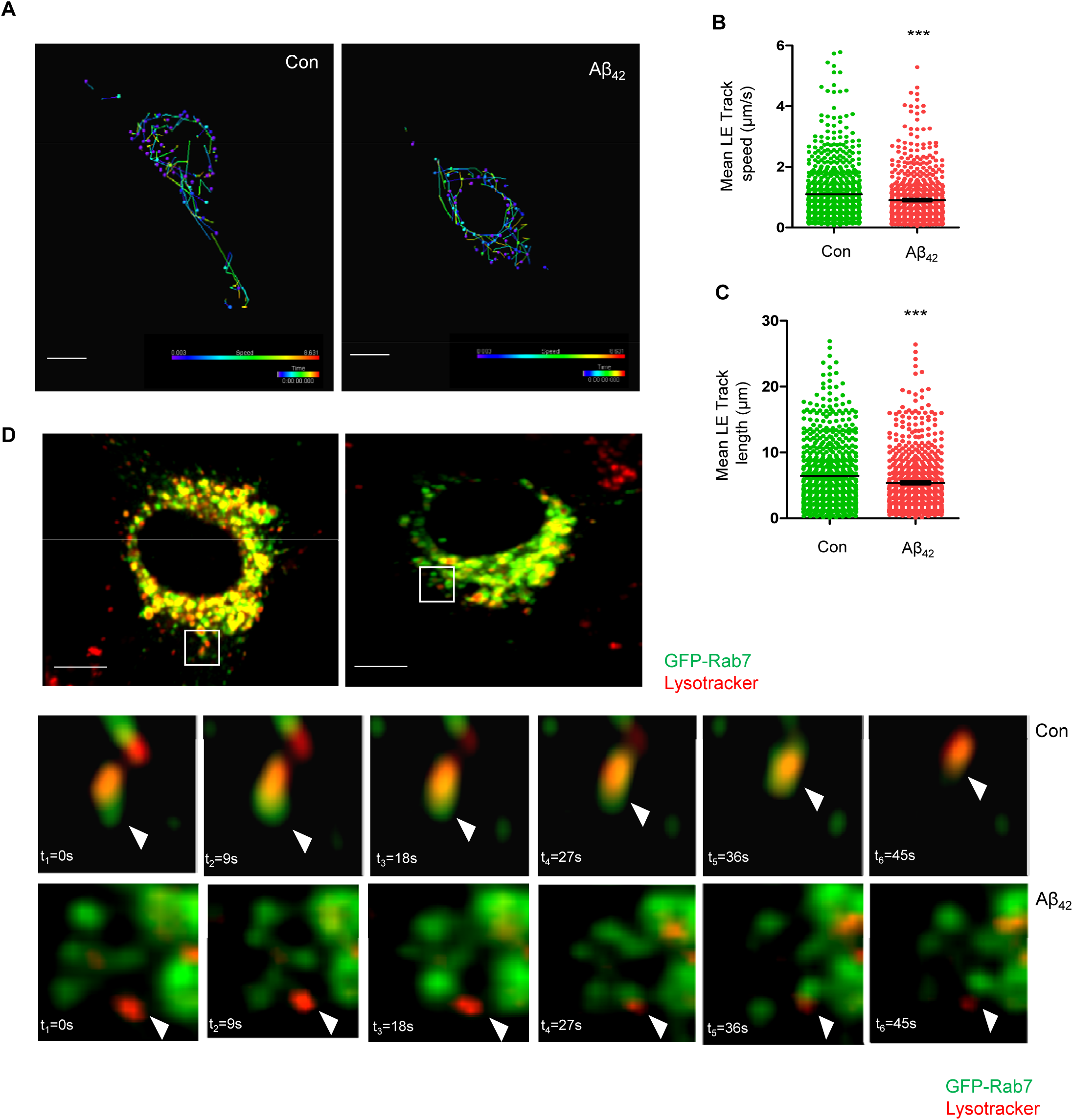
Effect of Aβ_1-42_ on Late Endosome dynamics. Graphs depicting mean track speed (B) and track length (C) of individual LEs in both control and Aβ_1-42_ treated cells. Representative images depicting movement of all the LEs taken from time lapse images (A). Track movement has been shown by white line and scale bar 10 μM. **D** Time lapse microscopic showing interaction between LE (green) and lysosome (red). C6 glioblastoma cells were transiently transfected with GFP-Rab7 (green) followed by 24 hrs of Aβ_1-42_ treatment. 100nM of lysotracker (red) was added in growing condition for 1hr before imaging. Time lapse images were taken with 9 sec gap for 90 seconds. The representative cells pictures are shown in upper panel with with marked regions. Time lapse images of these areas are shown in the panels below. For statistical significance, minimum three independent experiments were considered in each case unless otherwise mentioned and error bars are represented as mean ± S.E.M. P‐values were calculated by utilising Student’s t-test. ns: non‐significant, *P < 0.05, **P < 0.01, ***P < 0.0001. Scale bars 10 μm

**Supplementary Figure S4.**
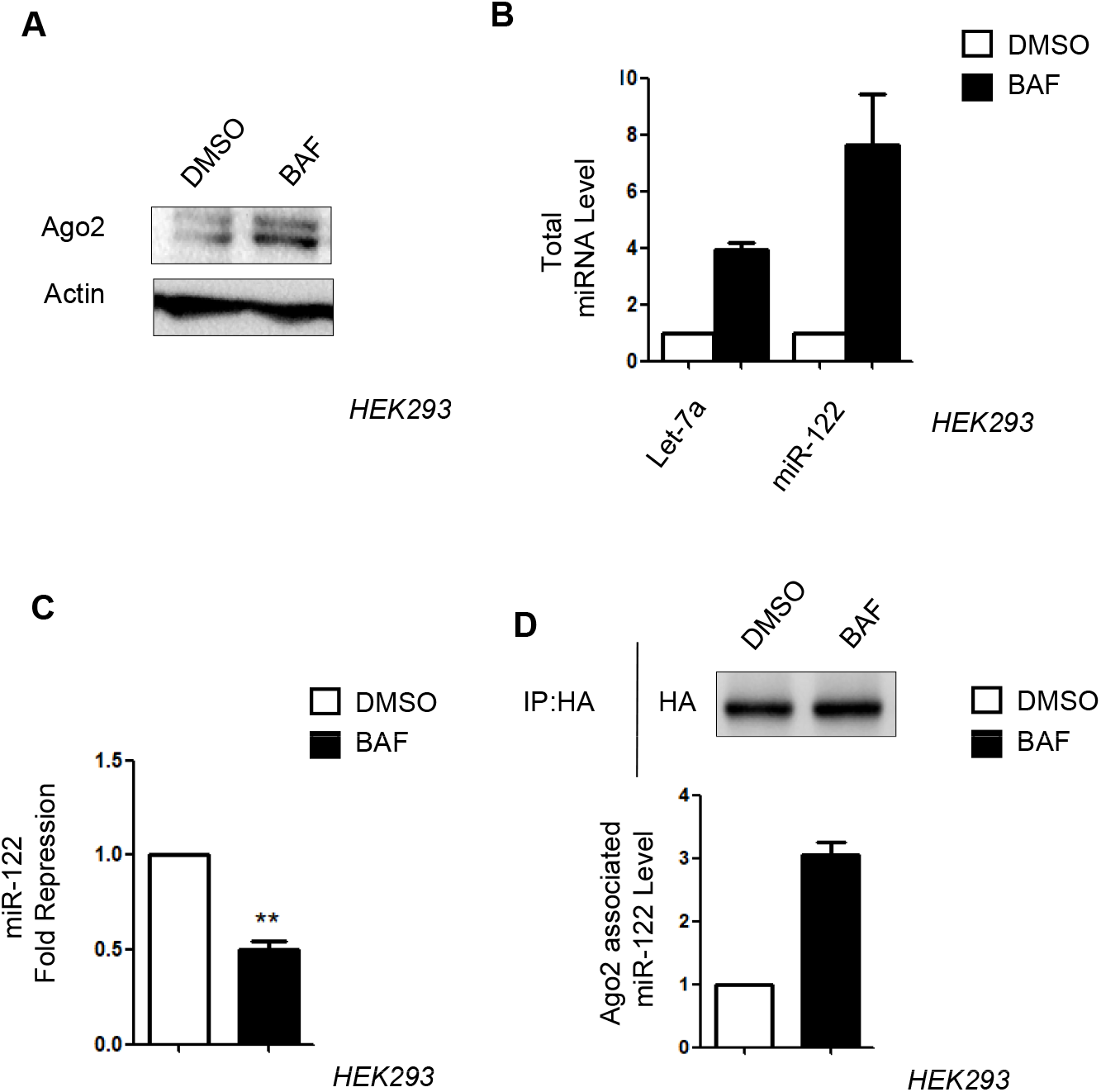
Effect of Bafilomycin on miRNA level and activity. **A** Western blot data showing cellular Ago2 level in HEK293 cells subjected to 100nM of Bafilomycin for 8 Hrs. **B** qPCR data showing cellular miRNA level upon bafilomycin treatment. qPCR data was normalized against U6 SnRNA. **C** Luciferase assay showing effect of bafilomycin (100nM for 8 Hrs.) on activity of miR-122. Pre-miR-122 plasmid along with its target RL-3xB-122 was exogenously transfected in HEK293 cells for this assay. **D** qPCR data showing amount of Ago2 associated miR-122 level upon bafilomycin treatment. HEK293 cells were transfected with Pre-miR-122 plasmid along with FLAG-HA-Ago2 plasmid. Immunoprecipitation followed by qPCR revealed the Ago2 bound miRNA level. qPCR data was normalized with amount of Ago2 recovered from IP experiment. For statistical significance, minimum three independent experiments were considered in each case unless otherwise mentioned and error bars are represented as mean ± S.E.M. P‐values were calculated by utilising Student’s t-test. ns: non‐significant, *P < 0.05, **P < 0.01, ***P < 0.0001.

**Supplementary Figure S5.**
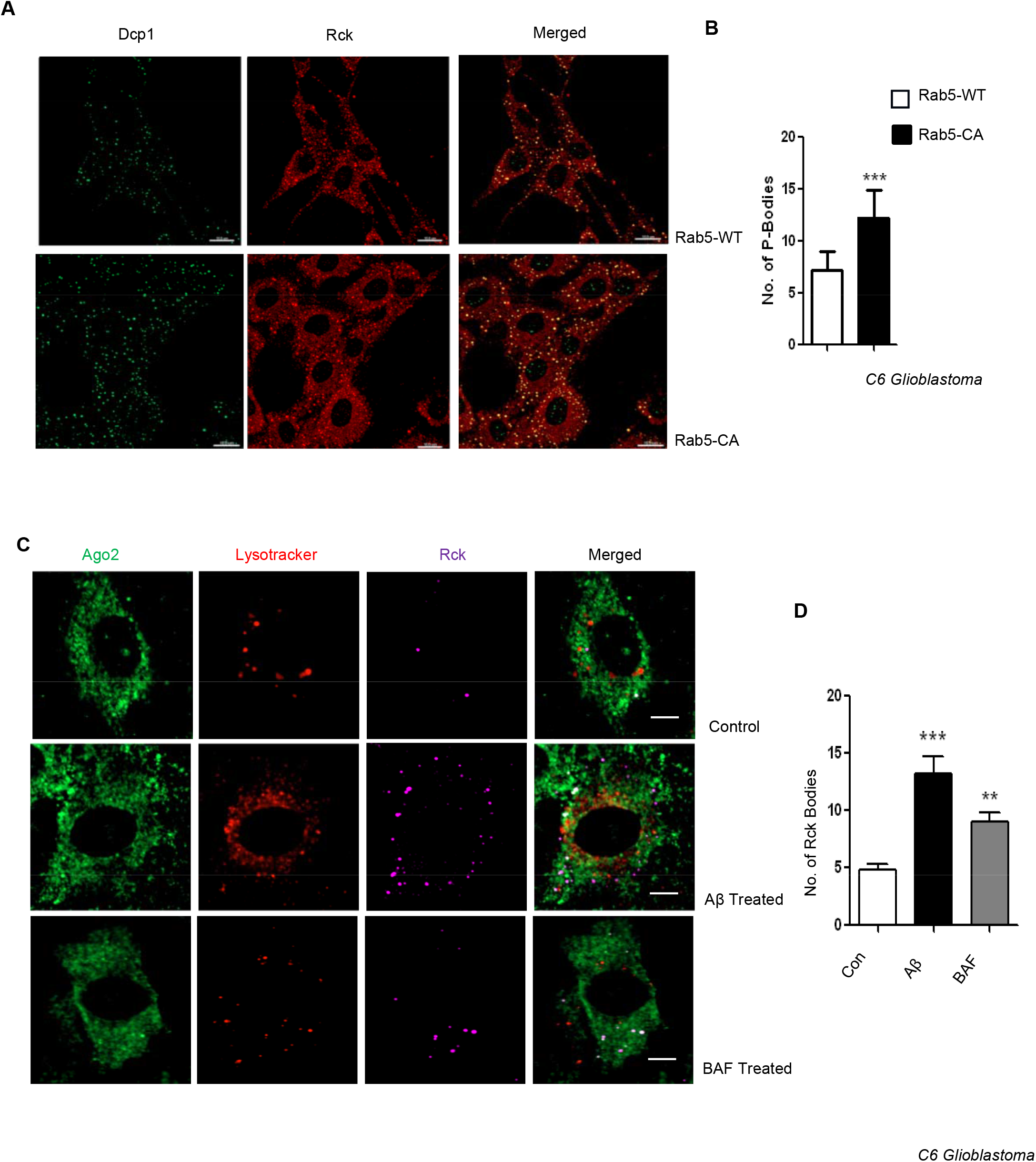
Perturbation of endosome maturation increases PB formation in mammalian cells. **A-B** Effect of expression of constitutively active Rab5-CA on PB formation in C6 glioblastoma cells. In confocal images of the cells Dcp1a (green) and Rck/p54 (red) were visualized by indirect immunofluorescence (A). Numbers of colocalized Dcp1a and Rck bodies were counted as P-bodies (B). **C-D** Effect of bafilomycin on PB formation in C6 glioblastoma cells. In confocal images Ago2 (green) and Rck/p54 (purple) were visualized by indirect immunofluorescence along with lysotracker (red) (C). Number of RCK/p54 bodies were counted as P-bodies after 8hrs of bafilomycin treatment (100nM) (D). For statistical significance, minimum three independent experiments were considered in each case unless otherwise mentioned and error bars are represented as mean ± S.E.M. P‐values were calculated by utilising Student’s t-test. ns: non‐significant, *P < 0.05, **P < 0.01, ***P < 0.0001. Scale bars 10 μm

**Supplementary Figure S6.**
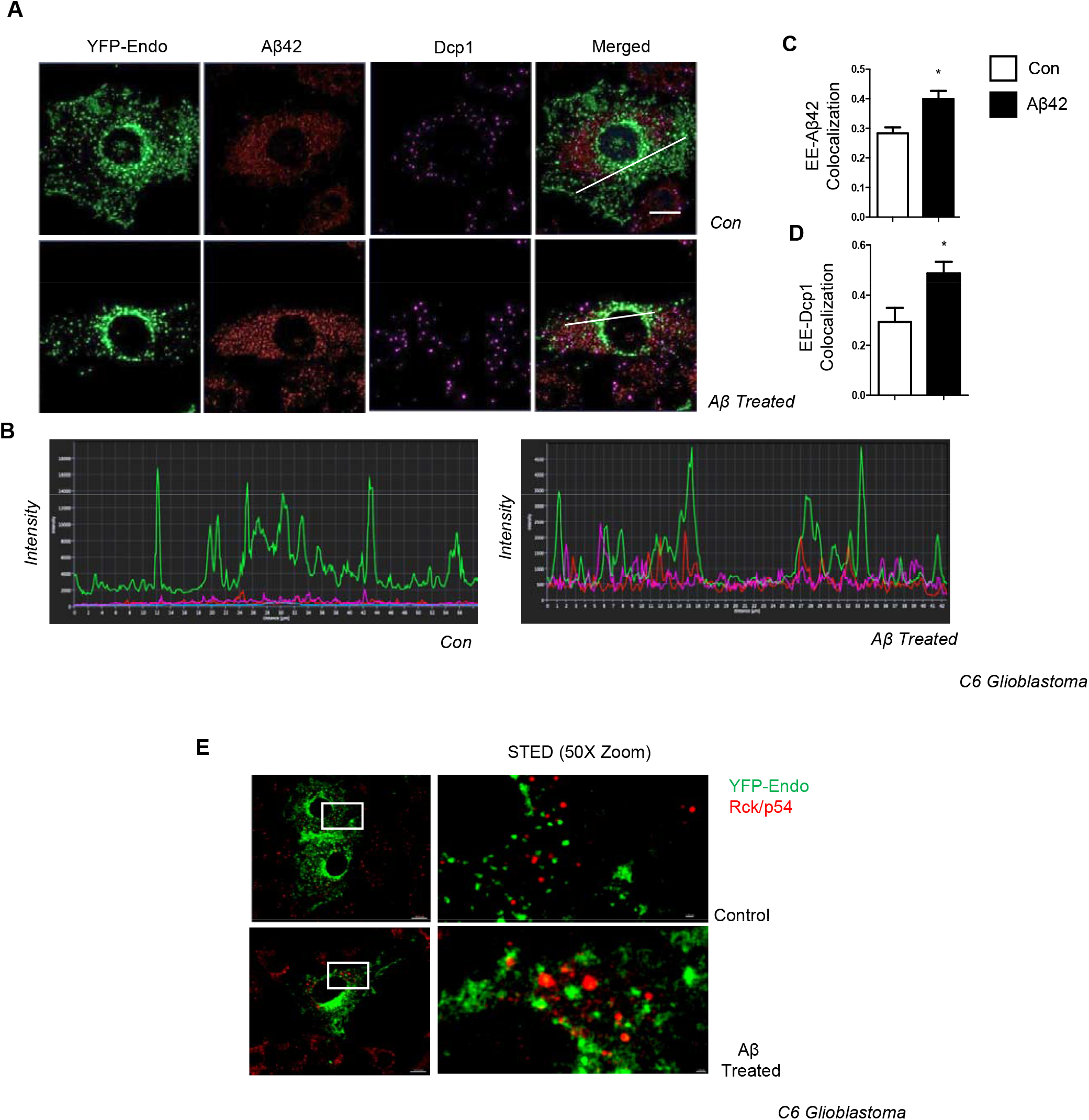
Juxtapositioning of PB components and endosomes after Aβ_1-42_ treatment of C6 glioblastoma cells. **A** Confocal images of the C6 Glioblastoma cells. YFP-Endo (green) was transiently transfected and Aβ_1-42_ (red), Dcp1 (purple) was visualized by indirect immunofluorescence in control and Aβ_1-42_ oligomer treated cells. **B** Localization and proximity of early endosome (EE) (green), Aβ_1-42_ deposition (red) and Dcp1 bodies (purple) in both control and Aβ_1-42_ treated cells. Graphs represents intensity profile of all the pixels along the defined section of the cells. **C-D** Graphs depicting Pearson’s coefficient of colocalization between both EE-Aβ_42_ and EE-Dcp1 in both DMSO and Aβ_42_ treated C6 cells. **E** Localization of early endosome (green) and PBs (red) in control and Aβ_1-42_ treated C6 Glioblastoma cells. YFP-Endo expressing cells were stained with endogenous Rck/p54 and observed under a STED microscope and zoomed part are shown. Scale bars 10 μm for non-zoomed part for Zoomed part, it is 1 μm. For statistical significance, minimum three independent experiments were considered in each case unless otherwise mentioned and error bars are represented as mean ± S.E.M. P‐values were calculated by utilising Student’s t-test. ns: non‐significant, *P < 0.05, **P < 0.01, ***P < 0.0001.

**Supplementary Table S1.**
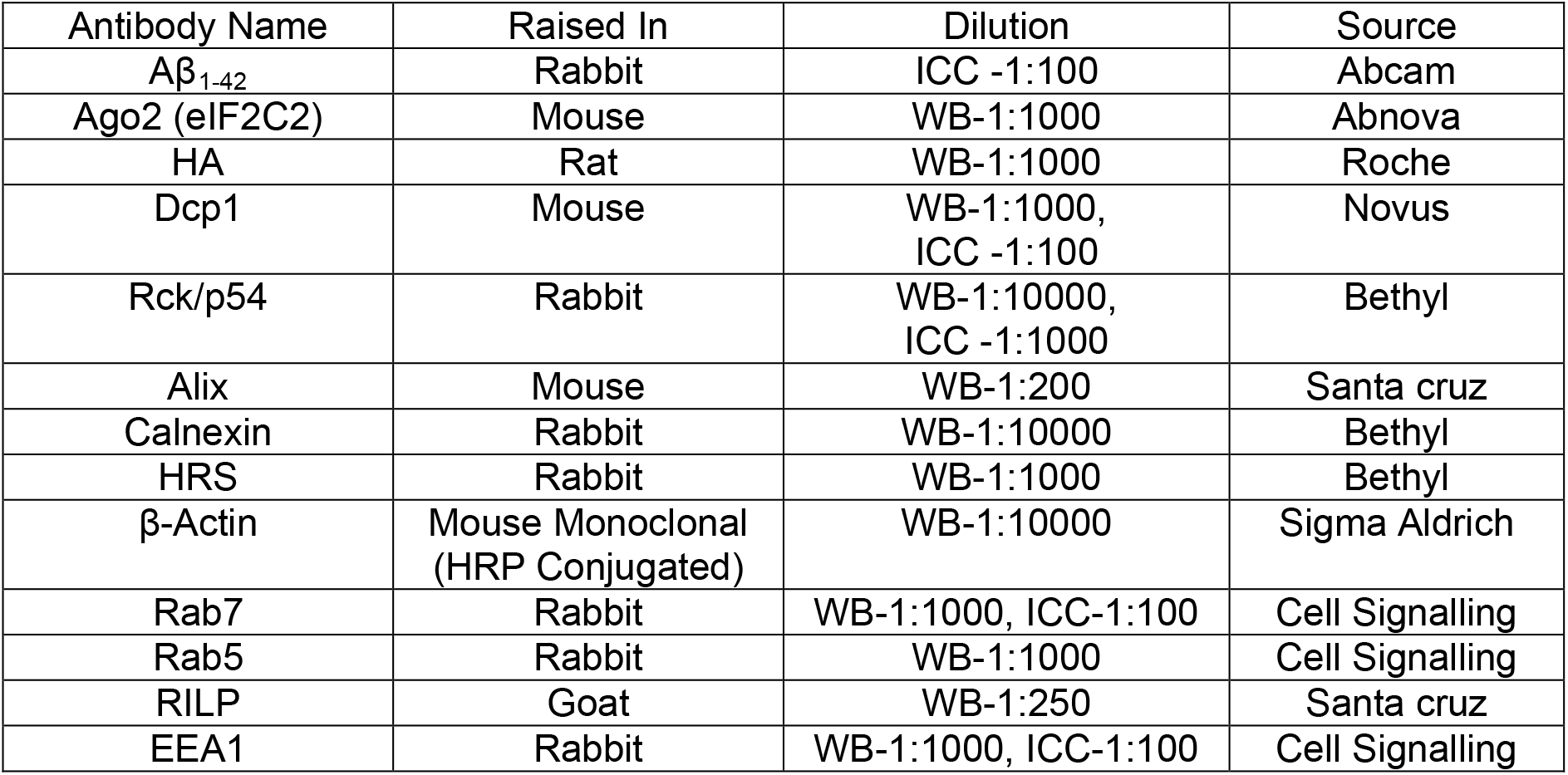
Details of Antibodies used for western blot (WB) and immunocytochemistry (ICC)

| Antibody Name | Raised In | Dilution | Source |
| --- | --- | --- | --- |
| Aβ<sub>1-42</sub> | Rabbit | ICC -1:100 | Abcam |
| Ago2 (eIF2C2) | Mouse | WB-1:1000 | Abnova |
| HA | Rat | WB-1:1000 | Roche |
| Dcp1 | Mouse | WB-1:1000,<br>ICC -1:100 | Novus |
| Rck/p54 | Rabbit | WB-1:10000,<br>ICC -1:1000 | Bethyl |
| Alix | Mouse | WB-1:200 | Santa cruz |
| Calnexin | Rabbit | WB-1:10000 | Bethyl |
| HRS | Rabbit | WB-1:1000 | Bethyl |
| β-Actin | Mouse Monoclonal<br>(HRP Conjugated) | WB-1:10000 | Sigma Aldrich |
| Rab7 | Rabbit | WB-1:1000, ICC-1:100 | Cell Signalling |
| Rab5 | Rabbit | WB-1:1000 | Cell Signalling |
| RILP | Goat | WB-1:250 | Santa cruz |
| EEA1 | Rabbit | WB-1:1000, ICC-1:100 | Cell Signalling |

**Supplementary Table S2.**
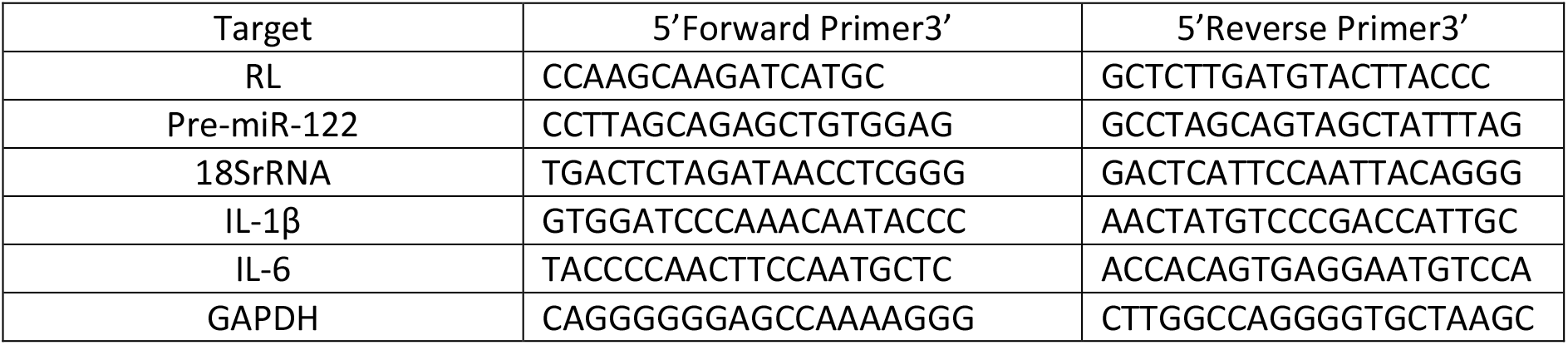
mRNA Primers used for quantification

**Supplementary Table S3.** Details of miRNA primers used for Taqman based quantification

| NAME | ASSAY ID |
| --- | --- |
| Let-7a | 000377 |
| miR-122 | 000445 |
| miR-146a | 000468 |
| miR-155 | 002571 |
| U6 SnRNA | 001973 |

